# Leaf Aqueous Extract of *Manilkara hexandra* influenced Glucose Metabolism in fish: An observation with *Labeo rohita* Fingerlings as model organism

**DOI:** 10.1101/2021.01.17.426992

**Authors:** Sumana Dutta, Debnarayan Chowdhury, Ria Das, Jayashri Das, Poulomi Ghosh, Koushik Ghosh, Sanjib Ray

## Abstract

*Manilkara hexandra* (Roxb.) Dubard (common name Khirni, family: Sapotaceae) is an evergreen traditionally used medicinal plant. The present study aimed to see the effects of leaf aqueous extract of *M. hexandra* (LAEMH) on digestive and glucose metabolic enzyme action and serum glucose level in rohu, *Labeo rohita*, fingerlings. Experimental fish were fed a basal diet (Group I), and diets supplemented with LAEMH at 300 mg (Group II) and 600 mg (Group III) kg^−1^body weight. A significant reduction in serum glucose was noticed in the treated groups when compared with the control group. The reduced amylase activity was noticed in the treated fingerlings in a dose-dependent manner. However, lipase and protease activities didn’t differ significantly among the experimental groups. Reduced serum glucose level might be correlated with the decline in the activity of digestive amylase in fish. Group II (3.35±0.19 U) and Group III (3.49±0.13 U) were recorded with reduced glucose-6-phosphatase activities than the control group (4.4±0.39 U). Moreover, the study revealed a decline in fructose-1,6-phosphatase activities in the treated groups in comparison to the control group. The decline in the activities of the metabolic enzymes might be associated with the non-availability of glucose owing to reduced activity of the digestive amylase in the treated groups. In conclusion, the present study established the hypoglycaemic effect of the leaf aqueous extract of *M. hexandra* in a fish model.

## 1. Introduction

*Manilkara hexandra* is an evergreen small to a medium-sized tree, 50-60 ft tall, and is distributed in the North, Central, and South India mainly in Gujrat, Madhya Pradesh, Maharashtra, Rajasthan, and tropical countries, also native to South Asia (Malik *et al*., 2012). The fruits, barks, and leaves are traditionally used for the treatment of opacity of the cornea, leprosy, ulcers, bronchitis, dyspepsia, urethrorrhea, *etc*. (Warrier *et al*., 1995; Chanda and Parekh, 2010). The different extracts of *M. hexandra* showed antibacterial, antiulcer, antioxidant, and antidiabetic activities (Mark *et al*., 2000; Dutta and Ray, 2020).

Earlier experiments on streptozotocin induced diabetic rats, it was observed that the leaf aqueous extract of *M. hexandra* could decrease fasting blood glucose time to time as compared to that of the metformin, an oral drug used to treat type 2 diabetes (noninsulin dependent diabetes mellitus) (Mark *et al*., 2000). Metformin has several effects like directly it inhibits complex I (NADH: ubiquinone oxidoreductase), AMP deaminase, glycerol-3-phosphate dehydrogenase and, in contrast activates AMPK via LKB 1(tumor suppressor gene), and all that leads to the inhibition of gluconeogenesis. Metformin also reduces the level of free fatty acids, triglycerides, LDL, lipid oxidation, and increase HDL level and thus reduces the risk of cardiovascular disease (DeFronzo and Goodman, 1995; Robinson *et al*., 1998; Landin *et al*., 1991).

Hyperglycemia and hyperlipidemia are two important factors that lead to the development of chronic diseases such as cardiovascular diseases and diabetes mellitus (Ogden *et al*., 2006). In the case of obese and overweight people, suppression of the postprandial lipid and glucose levels may reduce the occurrence of diabetes and its complications (Liao *et al*., 2006). So, the pancreatic lipase inhibitors and α-glucosidase inhibitors are used clinically for the treatment of hyperglycemia and hyperlipidemia (Van De Weel *et al*., 2005; Drew *et al*., 2007). In diabetic rats, hexokinase, glucose-6-phosphate dehydrogenase, and phosphoglucoisomerase activities were found to be at a decreased level, in contrast, glucose-6-phosphatase, and fructose1,6-bisphosphatase activities wherein increased level. But in treated diabetic rats the activity of hexokinase and phosphoglucoisomerase elevated and the level of glucose-6-phosphatase, fructose-1,6-bisphosphatase decreased indicating directly or indirectly inhibition of gluconeogenesis is an important strategy for antihyperglycemic therapy. There are several natural antihyperglycemic and antihyperlipidemic compounds that reduced carbohydrate and fat digestion or absorption (Önal *et al*., 2005; Seyedan *et al*., 2015). After administration of the different extract fractions showed recovery of hexokinase and glucose-6-phosphate dehydrogenase activity and decreased the glucose-6-phosphatase activity. For example, the bark extract of *Terminalia arjuna* treatment changed the activities of different enzymes of glycolysis and gluconeogenesis in rats. These effects of *T. arjuna* were due to the presence of flavonoids, tannin, and saponins (Ragavan and Krishnakumari, 2006). Administration of different extract fractions of the hydro-methanolic extract of *Swietenia mahagoni* seed to streptozotocin-induced diabetic male albino rats reduced the FBGL at different day’s intervals after the application. Among all the extract fractions, the n-hexane fraction of hydro-methanolic extract of *Swietenia mahagoni* showed effectiveness (Bera *et al*., 2015). The alkaloids like palmatine, jatrorrhizine, and magnoflorine isolated from *Tinospora cordifolia*, showed a significant reduction in hepatic gluconeogenesis (Patel and Mishra, 2011). Berberine, is an isoquinolone alkaloids isolated from *Coptis chinensis*, is a compound being used to treat type2 diabetes. It reduces the expression of PEPCK and G6Pase; inhibited mitochondrial function and also reduces FoxO1, SREBP1 transcription factors (Xia *et al*., 2011). Chlorogenic acid is a specific inhibitor of glucose-6-phosphate translocase in a rat microsome (Andrade-Cetto, 2010).

For antidiabetic activity screening of natural products, animals with abnormal glucose metabolism offer a good model system. There are several ways to develop rodents based diabetic models like - dietary/nutritional induction (Surwit *et al*., 1988), chemical induction (Dufrane *et al*., 2006), transgenic/knock-out manipulation (Butler *et al*., 2004), surgical manipulation (Risbud and Bhonde, 2002), a combination of the above (Srinivasan *et al*., 2005). But the major drawbacks of these techniques are ethical issues and labor-intensive process. Nowadays, zebrafish (*Danio rerio*) is being used widely as a model to study diabetes and various metabolic diseases as they possess physiological and metabolic similarities with mammals (Zang *et al*., 2017). Treatment of streptozotocin to zebrafish leads to the destruction of the adult pancreas and the blood glucose concentration increased up to 300 mg/dL (Olsen *et al*., 2010). Moreover, hyperglycaemic zebrafish and Indian perch developed retinopathy and increased glycosylated haemoglobin (Gleeson *et al*., 2007; Barma *et al*., 2006). Unlike herbivorous/omnivorous mammals, in these fish major energy for biological activities comes from protein metabolic pathways, gluconeogenesis, and TCA cycle. Fish is a good model system to study metabolic diseases and carps are generally known to be hyperglycemic. Therefore, the present study aimed to see the effects of leaf aqueous extract of *M. hexandra* on serum glucose level, digestive and glucose metabolic enzyme activities in *L. rohita* fingerlings.

## 2. Materials and methods

### 2.1. Chemicals

D-glucose, iodine, ammonium molybdate, and sulphuric acid was purchased from Qualigens Fisher scientific, Mumbai, India. Fetal bovine serum, NADP and FDP were obtained from Sigma-Aldrich, St. Louis, MO, USA. Tyrosine, DNSA, glucose-6-phosphate, pure olive oil, PVA, phenolphthalein indicator, ATP, MgSO_4_, MgCl_2_ and Tris HCl were purchased from Himedia Laboratories Pvt. Ltd., Mumbai, India. Folin phenol reagent, Starch, potassium dihydrogen phosphate, ANSA, TCA, 1-butanol, petroleum ether, methanol, and sodium hydroxide was obtained from Merck Specialities Pvt. Ltd., Mumbai, India. All other chemicals were of analytical grade and obtained from reputed manufacturers.

### 2.2 Plant material collection and aqueous extraction

Fresh leaves of *M. hexandra* were collected from the campus of The University of Burdwan.The herbarium specimen (No. BUGBSD015) is retained in the Department of Zoology for future reference. The collected leaves were washed in running tap water, shade dried, pulverized with an electric grinder, and stored in airtight container. Dried leaf powder was soaked with distilled water (1:10; w/v), boiled for 30 min in a water bath and the leaf aqueous extract of *M. hexandra* thus obtained was filtered through filter paper Whatman Filter paper 1 (150 mm). The leaf aqueous extract of *M. hexandra* (LAEMH) was stored at −20°C for further use.

### 2.3. Fingerlings culture and administration of LAEMH

*Labeo rohita* (Hamilton) fingerlings (average weight: 14.78±0.311 g) were collected from a local fish farm and acclimatized in fibre reinforced plastic (FRP) tanks (350 L) for 15 days before the experiment. The experimental fish were maintained and utilized for the study following the guidelines offered by the Dissection Monitoring Committee of The University of Burdwan. Post acclimatization, 126 healthy rohu fingerlings for 3 experimental groups were distributed in 9 FRP tanks (350 L) at a stocking density of 14 fish per tank with three replicates for each group. The basal diet (consisting of fish meal 30%, mustard oil cake 20%, and rice bran 50%) without LAEMH supplementation was fed to the control group (Group I), whereas basal diet was fortified with the dried LAEMH so as to feed the experimental groups at 300 mg (Group-II) and 600 mg (Group-III) per kg body weight of fish. Chromic oxide (1%) was added separately as an external digestibility marker. The fish were fed twice daily: at 08.00 h and 14.00 h, at a feeding rate of 3% (w/w) of the total body weight per day and the feeding trial was continued for 60 days. Level of incorporation of LAEMH was calculated on the basis of pooled weight of fish in a group at the specified feeding rate, and LAEMH supplemented diets were prepared at every 10^th^ day interval to ensure required concentration of LAEMH in the diet. Pelleted diets were dried in a hot-air cabinet (60^0^C), packed in sterile airtight polythene bags and stored at 4°C in a refrigerator. The faecal samples released by the fish were collected from each aquarium following the “immediate pipetting” method outlined by Spyridakis *et al*. (1989). The oven dried (60°C) faecal samples were analysed for digestibility estimation. Water quality parameters (e.g., temperature 25-28°C, pH7.1-7.4 and dissolved oxygen 6.7-7.3 mg L^−1^) were monitored after American Public Health Association (APHA, 2012) and varied within a narrow range during the experimental period.

### 2.5. Digestive enzymes assays

Activities of the digestive enzymes were analyzed in triplicate, prior to commencement and at the termination of the experiment. For each of the replicates, three fish from each tank were sampled, anesthetized by applying 0.03% tricaine methane sulfonate (MS-222), gastrointestinal (GI) tracts were removed and placed in chilled Petri plates. Blood, along with mesenteries and debris were cleaned with chilled phosphate-buffered saline (PBS; 0.1M, pH 7.4, 0.89% sodium chloride). A 10% homogenate in PBS was prepared using a tissue homogenizer (REMI, Model RQ-127A2), centrifuged (10,000*g*, 20 min, 4^0^) and the supernatant was evaluated for determination of the enzyme activity. The protein content of the supernatant was estimated following Lowry *et al*. (1951) using bovine serum albumin as standard. Protease activity was determined using *Hammerstein* case in (substrate) after Walter (1984). Enzyme activity (unit) was defined as µg of tyrosine liberated mg protein^−1^ h^−1^. Amylase (α-amylase) activity was measured following Bernfeld (1955) using a dinitro-salicylic-acid (DNSA) reagent. Amylase activity was expressed as mg maltose liberated mg protein^−1^ h^−1^. Lipase activity was determined according to the method described by Bier (1955) using olive oil as substrate. Lipase activity was expressed as µ mole of fatty acid liberated mg protein^−1^ h^−1^.

### 2.6. Glucose metabolic enzymes assays

After collecting the GI tracts for analyses of the digestive enzymes, hepato-pancreatic tissues were removed and collected separately (in chilled 0.25 M sucrose solution, pH 7.4) for preparation of samples for metabolic enzymes, a detailed description of which was presented in Mondal *et al*. (2020). The mitochondrial and cytosolic tissue fractions were kept at −20°C until use.

Glucokinase activity was measured after Traulis *et al*., (1996) using glucose as a substrate. The enzyme activity unit was defined as the amount of enzyme that phosphorylates 1 µM of glucose min^−1^ under assay condition. Glucose-6-phosphatase (G-6-Pase) activity was measured following Marjorie (1964) using glucose-6-phosphate (0.1 M) as substrate and malate buffer (0.1 M, pH 7.4). Fructose-1, 6-bisPhosphatase (F-1,6-BPase) activity was determined after Freeland & Harper (1959) that utilized fructose-di-phosphate (0.1M, pH 7.4) as substrate in borate buffer (0.1 M, pH 7.4). Release of phosphate (Pi) in both the methods was estimated following Fiske & Subbarow (1925), and enzyme activities were expressed as µg of phosphorus released mg^−1^ protein min^−1^. Glucose-6-phosphate dehydrogenase (G6PD) activity was determined according to Kornberg & Horecker (1955) using glucose-6-phosphate (substrate) and NADP in Tris-HCl buffer (0.08 M, pH 7.4). Enzyme activity was expressed as µM of NADPH formed mg^−1^ protein h^−1^.

### 2.7. Estimation of serum biochemical parameters

Serum was collected from fish blood following Ai *et al*. (2006). Fish were starved on the previous day (24 h), anesthetized and blood samples were collected from the caudal vein (Khan *et al*. 2015). For each replicate, a pooled sample from three fish was allowed to clot at room temperature (25^0^C, 4 h). Afterward, the sample was centrifuged (836*g*, 10 min, 4°C), serum was removed and stored at −20 °C until use.

The serum biochemical parameters like triglyceride, cholesterol, protein, albumin, alkaline phosphatase, glucose, HDL, LDL, SGPT, etc. were estimated from the blood serum of aqueous extract treated and untreated fish through the biochemical analyzer Erba Mannheim (EM DESTINY 180) using commercial kits following manufacturer’s instruction.

### 2.8. Determination of apparent digestibility

Digestibility is the quantification of the digestive processes. It provides a relative measure of the extent to which ingested food and its nutrient compounds have been digested by the fish. Two types of digestibility estimation were conducted for rohu fingerlings. Chromic oxide levels in the diets and fecal samples were estimated spectrophotometrically following Bolin *et al*. (1952) involving perchloric acid digestion. Dry matter and other nutrient levels were estimated in chromic oxide incorporated diets as well as the fecal samples following the methods of AOAC (1990) as described earlier. Apparent dry matter or total and nutrient digestibility values calculated employing the following formulae (De Silva and Anderson, 1995).

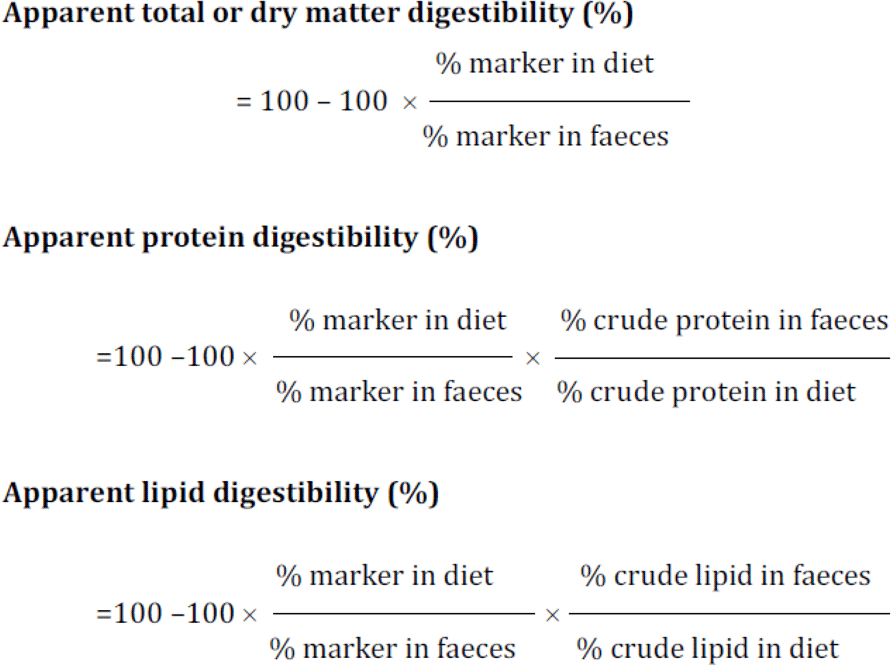

### 2.9. Statistical analysis

All the assays were performed at least in triplicate and all the data points were expressed as Mean**±**SEM. The statistical significance of differences between the groups was determined by two population Student’s t-Test **(\***at *p*< 0.05, **\*\***at*p*< 0.01, ***at *p*< 0.001) and One-Way Analysis of Variance (ANOVA) (**Significant at *p*< 0.01). Differences between means were regarded as significant at *p*< 0.05 (Tukey test). The analysis was performed by using Origin Pro software 2019 (Academic) 32-bit.

## 3. Results

### 3.1. Effect of LAEMH on the digestive enzymes

After two months of LAEMH oral administration with food on *Labeo rohita* fingerlings, there was a significant reduction in amylase activity in the treated groups as compared to the control group. Here, a reduced amylase activity (14.03±0.49,13.68±1.19 and 10.72±3.61µg maltose released mg protein ^−1^h^−1^ (*p*<0.05) respectively at 0,300 and 600 mg/kg body weight) was observed on LAEMH administration on *Labeo rohita* fingerlings. In the case of protease activity, it shows an increased (8.86±0.161 to 9.11±0.64µg tyrosine liberated mg protein ^−1^h^1^) tendency in Group III. But in the case of Group II, the protease activity slightly decreased from the control group though the differences are not statistically significant. The differences in lipase activities (9.063±0.33U, 9.56±0.18 U, and 9.14 ±0.77 U respectively for Group I, Group II, and Group III) are not statistically significant (Table 1).

**Table 1.**
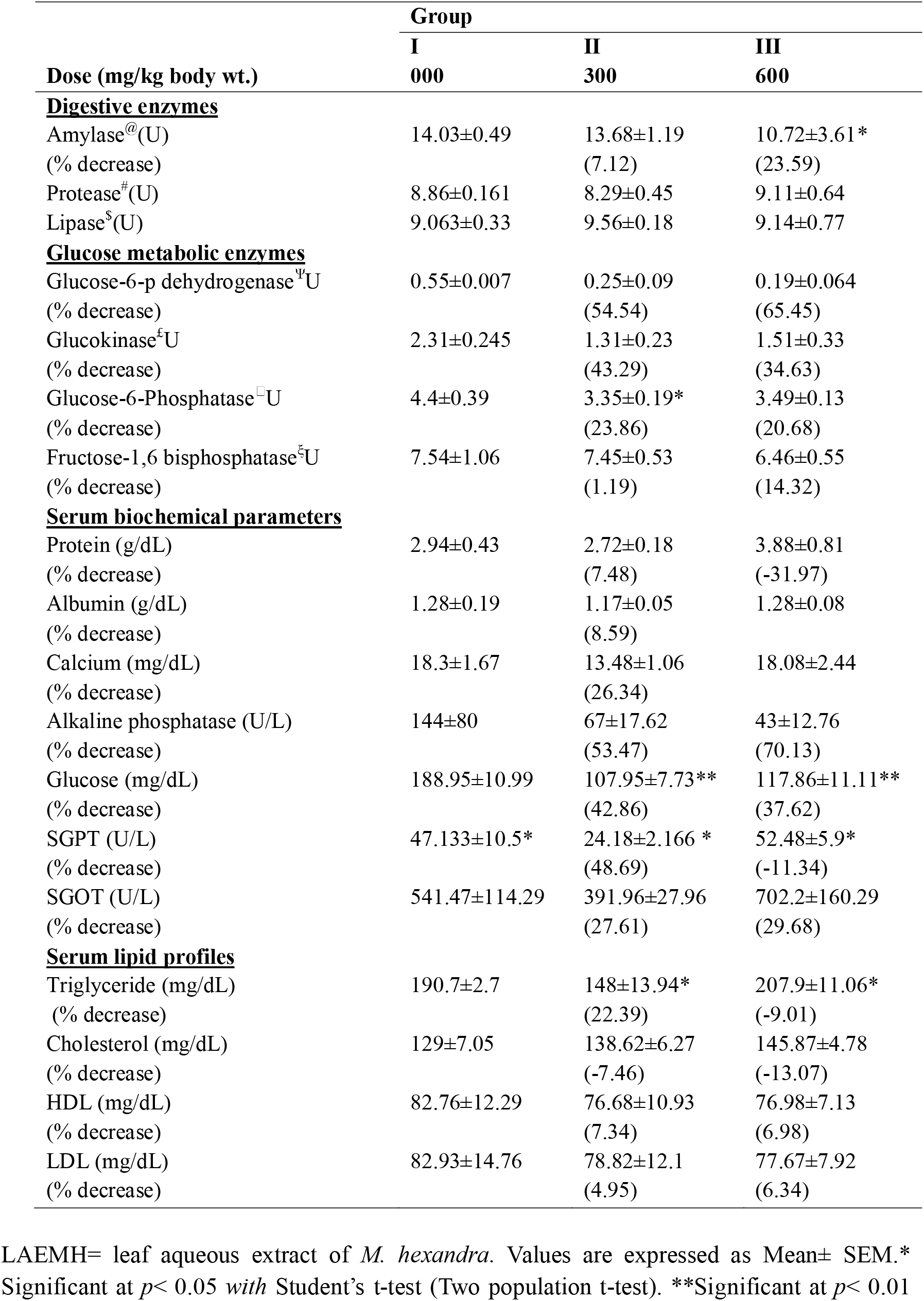
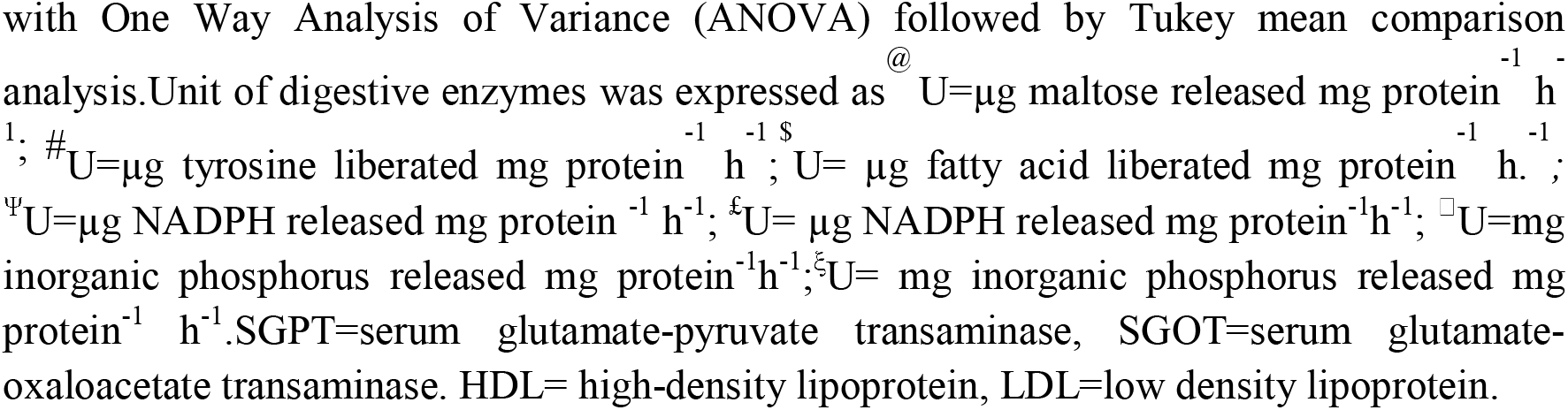
Effect of LAEMH on digestive and glucose metabolic enzymes, serum biochemical and lipid profiles of *Labeorohita*aftertwo months of continuous treatment

### 3.2. Effect of LAEMH on the glucose metabolic enzymes

Treatment of LAEMH on *Labeo rohita* fingerlings showed a dose-dependent decrease of glucose-6-phosphate dehydrogenase activity. In the control group, Group I, G6PD activity was recorded as 0.55±0.007 U whereas that was found be decreased in case of LAEMH treated and the enzyme activity values were 0.25±0.09 U (54.54% decrease) and 0.19±0.064U (65.45% decrease) respectively for Group II and Group III. Like G6PD, glucose-6-phosphatase the activity was also found to be decreased in LAEMH treated groups; Group II (3.35±0.19U) and group III (3.49±0.13 U) than the control group (4.4±0.39 U, µg NADPH acid liberated mg protein ^−1^h^−1^). The LAEMH treatment also reduced the activity of fructose-1,6-phosphatase intreated groups (14.32% fructose-1,6-phosphatase activity reduced in Group III after 2 months of the experiment than the control Group I.

### 3.3. Effect of LAEMH on the serum biochemical parameters

The serum glucose concentration of *Labeo rohita* fingerlings was significantly decreased in the case of the treated groups (Group II: 107.95±7.73 mg/dL; reduced 42.86% and Group III: 117.86±11.11 mg/dL; reduced 37.62%) as compared to that of the untreated group (Group I: 188.95±10.99 mg/dL) after 2 months of treatment (Table.1). The serum protein level decreased in group II (2.72±0.18 U) but increased in Group III (3.88±0.81U) as compared to the Group I (3.88±0.81U). In the case of serum albumin level, the values are the same in three groups but it is slightly decreased in Group II (1.17±0.05 U) as compared to Group I (1.28±0.19 U) and II (1.28±0.08 U). Serum calcium level also decreased (26.34 %) in Group II but Group III fish maintained the same level to that of the control Group I. There was a dose-dependent decrease in the serum alkaline phosphatase level. Both serum SGPT and SGOT levels were decreased in group II as compared to the control. The serum SGPT level was decreased from 47.133±10.5 to 24.18±2.166 U/L in the case of Group II. The serum SGOT level also decreased from 541.47±114.29 to 391.96±27.96 U/L in the case of Group II. But both these levels significantly increased in Group III (52.48±5.9 and 702.2±160.29 U/L respectively). In the case of serum lipid concentration, the triglyceride level decreased in Group II (148±13.94 mg/dL) but it was increased in Group III (207.9±11.06 mg/dL) as compared to that of the control group (145.87±4.78). Whereas the serum cholesterol level gradually increased with the doses as compared to group I. Both HDL and LDL levels decreased in treated groups than the untreated groups but the differences were not statistically significant (Table 1).

### 3.4. Effect of LAEMH on the Growth performances of *Labeo rohita* fingerlings

Data regarding growth performance and utilization of nutrients by *L. rohita* fingerlings fed experimental diets are presented in Table 2. The average final weight of the fish increased considerably from the initial value in all the groups but the final weight of the treated group does not differ as compared to the control (Group I) group. The highest value (2.62 ± 0.04) of FCR for Group III was noticed in fish fed LAEMH 600 mg/kg body wt. The lowest value of FCR (2.44 ± 0.03) was observed in Group I. The value of SGR was the lowest in fish treated 600 mg/kg body wt LAEMH whereas the highest PER (1.36 ± 0.03) was observed in Group I.

**Table 2.** Growth performances, apparent digestibility of dry matter and nutrients of *Labeo rohita* fingerlings fed experimental diets for 60 days.

| Dose (mg/kg body wt.) | Group |  |  |
| --- | --- | --- | --- |
|  | I<br>000 | II<br>300 | III<br>600 |
| **Growth performances** |  |  |  |
| Initial Weight (g) | 15.00 ± 0.55 | 14.99 ± 0.42 | 14.39 ± 0.47 |
| Final Weight (g) | 31.54 ± 0.67 | 30.47 ± 0.54 | 28.17 ± 0.58 |
| Feed Conversion Ratio (FCR) | 2.44 ± 0.03 | 2.48 ± 0.04 | 2.62 ± 0.04 |
| Specific Growth Rate (SGR) | 1.23 ± 0.03 | 1.18 ± 0.02 | 1.11 ± 0.02 |
| Protein Efficiency Ratio (PER) | 1.36 ± 0.03 | 1.34 ± 0.03 | 1.27 ± 0.02 |
| **Apparent digestibility of dry matter and nutrients** |  |  |  |
| Dry matter % | 67.6 ± 0.62 | 62.85 ± 0.87 | 61.67 ± 0.52 |
| Protein % | 86.49 ± 0.70 | 84.42 ± 0.67 | 86.74 ± 0.95 |
| Lipid % | 85.03 ± 0.71 | 85.42 ± 0.86 | 83.46 ± 0.67 |

### 3.5. Effect of LAEMH on the apparent dry matter and nutrient digestibility of *Labeo rohita* fingerlings

The highest apparent dry matter digestibility (ADD) was observed in Gr. I. although there was no significant difference in its values as compared to the treated groups. There was no significant difference in apparent protein and lipid digestibility (APD & ALP) among the different groups (Table 2).

## 4. Discussion

Ethno-medicinal plants play a major role in the treatment of prolonged disease, diabetes mellitus, especially in developing countries. Diabetes mellitus is accompanied by several other chronic disorders such as polyuria, polydipsia, glucosuria, *etc*. The treatment of each of such diseases can be done by exploiting the herbal integrity of India. Research on natural products plays an important role in discovering and developing new drugs, which are having hopefully more effective and no side actions compared to most modern drugs. For antidiabetic activity screening of natural products, animals with abnormal glucose metabolism offer a good model system. Glucose tolerance tests in fish resulted in prolonged hyperglycemia like the diabetic state in mammals (Hertz *et al*., 1989). Low sensitivity of insulin towards glucose levels in fish has been suggested (Hertz *et al*., 1989). Therefore, the present study utilized fish (an Indian major carp, *Labeo rohita*) as a model to appraise the anti-diabetic effects of leaf aqueous extract of *Manilkara hexandra* through analyzing its effects on amylase, protease, lipase, glucokinase, glucose-6-phosphatase (G-6-Pase), fructose-1,6-bisphosphatase (F-1,6BPase), Glucose-6-phosphate dehydrogenase (G6PD), alkaline phosphatase, serum glucose, serum glutamic oxaloacetic transaminase (SGOT), serum glutamic pyruvic transaminase (SGPT), serum total protein, cholesterol, low-density lipoprotein (LDL), and high-density lipoprotein (HDL).The decline in amylase activity in fish fed LAEMH incorporated diets was noticed. However, protease and lipase activities didn’t differ significantly among the experimental groups. Non-specific inhibition of digestive enzymes in fish exposed to plant extracts or plant ingredients in the diets have been reported previously (Mandal and Ghosh, 2018). However, inhibition of only amylase activity among the studied digestive enzymes could be due to specific inhibition caused by LAEMH. Leaf methanolic extract of *M. hexandra* showed α-amylase inhibitory activity both *in vivo* and *in vitro*. Ethyl acetate fraction of leaf methanolic extract of *M. hexandra* showed the highest α-amylase inhibitory activity in streptozotocin-induced diabetic rats as compared to the standard drug acarbose (Patel and Patel, 2015). Thus, our study is in agreement with the previous report showing the α-amylase inhibitory property of the extract from *M. hexandra* leaves (Patel and Patel, 2015).

In the present study, reduced activities of the enzymes for carbohydrate metabolism were noticed in fish fed diets with LAEMH. Hexokinase/glucokinase is the key enzyme in the glycolytic pathway that catalyzes the conversion of glucose to glucose-6-phosphate and indicates glucose utilization by the glycolytic pathway. We observed here declined glucokinase activities in fish treated with increasing dose of LAEMH incorporation in the diets, although differences were not significant (*p*<0.05).

Glucose-6-phosphatase (G6Pase) is another important target for type 2 diabetes as its expression is increased in type 2 diabetes. G6Pase is the enzyme of the final reaction of of gluconeogenesis (Westergaard *et al*., 2002). Targeting the catalytic site of the enzyme is very effective in antihyperglycemic therapy. But there are some limitations also like severe hypoglycemia and fatty liver may be the side effects (Agius, 2007). The other important strategies are like inhibition of phosphoenolpyruvate kinase (PEPCK) and fructose-1,6-bisphosphatase (F1,6BPase). But inhibition of F-1,6BPase is effective in mild but not in severe diabetes (Agius, 2007). In the present study, glucose-6-phosphatase (G-6-Pase) and fructose-1,6-bisphosphatase (F-1,6BPase) activities were evaluated that help to produce glucose from non-carbohydrate precursors (amino acid, lactate, glycerols, *etc*.) for glucose homeostasis. Significantly lower (*p*<0.05) G-6-Pase activity in fish fed 300 mg LAEMH kg^−1^ body weight was noticed in the present study, however, F-1,6BPase activity didn’t differ significantly. Reduction of glucose-6-phosphatase activity was also observed in *L. rohita* after clove and cardamom treatment (Asimi and Sahu, 2016). Inhibition of G-6-Pase activity was also reported when *L. rohita* was exposed to phenolic compounds; viz., monohydric phenol, and m-cresol, which might suggest microsomal membrane damage as the enzyme was known to be located exclusively in the endoplasmic reticulum membrane (Gaur *et al*., 2019). In the case of treated diabetic rats, the activities of glucose-6-phosphatase and fructose-1,6-bisphosphatase were decreased. These effects of *T. arjuna* were due to the presence of flavonoids, tannin, and saponins (Ragavan and Krishnakumari, 2006). The reduction in the enzyme activity might be indicative of a decrease in *de novo* enzyme protein synthesis or substrate deficiency.

Glucose-6-phosphate dehydrogenase (G6PD) is directly involved in lipogenesis and stress management by providing NADPH through the HMP-shunt (pentose phosphate pathway). Glucose-6-phosphate dehydrogenase is a regulatory protein that produces reducing equivalents for fatty acid biosynthesis and the recycling of pentose sugars in energy metabolism (Beutler, 1991). In this study, liver G6PD activity was noticed to be declined in fish fed LAEMH incorporated diets, although not significantly. There are several plant-derived compounds that inhibit gluconeogenesis either by inhibiting enzymes of this pathway or by activating AMPK. Thus, directly or indirectly inhibition of gluconeogenesis is an important strategy for antihyperglycemic therapy. Chlorogenic acid is a specific inhibitor of glucose-6-phosphate translocase in a rat microsome (Andrade-Cetto, 2010). The alkaloids like palmatine, jatrorrhizine, and magnoflorine isolated from *Tinospora cordifolia*, showed a significant reduction in hepatic gluconeogenesis (Patel and Mishra, 2011).

Serum biochemical parameters are considered important tools to examine the metabolic and physiological status of fish. In the present study, the hypoglycemic effect of LAEMH became evident as a significantly lower serum glucose level was recorded with the application of LAEMH in the diets. Oral administration of 50% bark ethanolic extract of *M. hexandra* (250 & 500 mg/kg body wt.) for 21 days to diabetic rats showed a reduction in serum glucose level. Administration of the methanolic bark extract shows the hypoglycemic effect on both diabetic and normal rats reduced the blood glucose at different time intervals (Nimbekar *et al*., 2012). Ethyl acetate fraction of leaf methanolic extract of *M. hexandra* showed the highest inhibitory activity among all the extract fractions (Patel and Patel, 2015). In alloxan-induced diabetic rats, ethyl acetate fraction of *Pterocurpus marsupium* efficiently lowers the blood glucose level after treated with this fraction (Ahmad *et al*., 1991). The treatment of ethanolic leaf extract of the *Croton zambesicus* in alloxan-induced diabetic rats reduced the blood glucose level that is comparable to that of the standard drug chlorpropamide (Okokon *et al*., 2006). A prominent decrease in blood glucose concentration of rohu, after exposure to phenolic compounds (phenol and m-cresol), was reported previously (Gaur *et al*., 2019). Hori *et al*., (2006) reported an insignificant decrease (*p*<0.05) in the plasma glucose levels of Red-tailed Brycon, *Bryconcephalus* after 96 h exposure to phenol. Several phytochemicals were noticed to be present in the LAEMH, e.g., tannin, phlobatannin, flavonoids, alkaloids, carbohydrates, saponin, and glycosides (Dutta and Ray 2016). A wide variety of phytochemicals were isolated and identified from time to time by different groups of workers from the different parts of *M. haxandra*. From leaves, hentriacontane, a triterpene ketone, cinnamic acid, a terpenic hydrocarbon, besides quercitol and taraxerol were isolated (Misra and Mitra, 1968). Moreover, tannins and terpenoids were present abundantly in comparison to the other compounds. Tannins are also phenolic derivatives. It has been reported that tannin can inhibit amylase activity even at very low concentrations. Tannin isolated from *Acacia auriculiformis*reduced the activities of amylase, protease, lipase isolated from *L. rohita*(Hamilton) fingerlings (Maitra and Ray, 2003). Tannin from *Pistia* leaf also inhibited digestive enzymes of *L. rohita* fingerlings in a dose-dependent manner (Mandal and Ghosh, 2010). Thus, inhibition of amylase activity by LAEMH might be due to the presence of a high amounts of tannins or other phenolics present in the extract.

Serum SGOT and SGPT activities decreased with the application of 300 mg LAEMH kg^−1^ body weight, while increased with 600 mg LAEMH kg^−1^ bodyweight. Thus, a higher dose of LAEMH (600 mg kg^−1^ body weight) might be indicative of dietary stress in fish. Exposures to phenolic compounds were also reported to portray an increasing trend of SGOT and SGPT in rohu (Gaur *et al*., 2019) and rainbow trout (Monfared and Salati, 2012). Further, serum total protein, cholesterol, low-density lipoprotein (LDL), and high-density lipoprotein (HDL) were remained more or less unaffected due to dietary application of LAEMH in fish.

In summary, activities of the enzymes related to carbohydrate metabolism were found in decreased level in the LAEMH treated fingerlings. Glucose-6-phosphatase reduced significantly in low doses as compared to the control group. The study recorded a decline in the activities of the key enzymes involved in glycolysis, gluconeogenesis, or HMP-shunt. Thus, it appeared likely that LAEMH-induced hypoglycemia was not due to increased utilization of glucose by the metabolic processes. On the contrary, the non-availability of glucose might have resulted in owing to a decline in the activity of digestive amylase.

## Data availability Statement

Research data are not shared. The data available from the authors upon reasonable request.

## Acknowledgment

The authors acknowledge the financial support of the DST INSPIRE fellowship (IF 110690) under INSPIRE program and infrastructural facilities (UGC-MRP, DST-FIST, DST-PURSE, and UGC-DRS sponsored) of the Department of Zoology, The University of Burdwan, West Bengal, India.

## References

Agius, L. (2007). New hepatic targets for glycaemic control in diabetes. Best Practice & Research Clinical Endocrinology & Metabolism, 21, 587–605.

Ahmad, F., Khalid, P., Khan, M.M., Chaubey, M., Rastogi, K., & Kidwai, J.R.(1991). Hypoglycemic activity of Pterocarpus marsupium wood. Journal of Ethnopharmacology,35(1), 71–75.

Andrade-Cetto, A. (2011). Inhibition of gluconeogenesis by Malmeadepressaroot. Journal Of Ethnopharmacology, 137, 930–933.

Andrade-Cetto, A., & Wiedenfeld, H. (2004). Hypoglycemic effect of Acosmiumpanamensebark on streptozotocin diabetic rats. Journal of Ethnopharmacology, 90, 217–220.

Asimi, O.A., & Sahu, N.P. (2016). Effect of antioxidant rich spices, clove and cardamom extracts on the metabolic enzyme activity of Labeorohita. Journal of Fisheries and Livestock Production, 4(1), 1–6.

Barma, P., Dey, D., Basu, D., Roy, S.S., & Bhattacharya, S. (2006). Nutritionally induced insulin resistance in an Indian perch: a possible model for type 2 diabetes. Current Science, 90, 188–194.

Bera, T.K., Chatterjee, K., & Ghosh, D. (2015). Remedial hypoglycemic activity of n-hexane fraction of hydro-methanol extract of seed of Swietenia mahagoni (L.) Jacq. in Streptozotocin-induced diabetic rat: A comparative evaluation. Journal of Herbs, Spices & Medicinal Plants, 21, 38–58.

Bernfeld, P. (1955). Amylase (alpha) and (beta). In: Colowick SP and Kaplan NO (Eds). Methods in enzymology (pp. 149–150). Academic Press, New York.

Beutler, E. (1991). Glucose-6-phosphate dehydrogenase deficiency.The New England Journal of Medicine, 324(3), 169–174.

Bier, M. (1955). Lipases. In: Colowick SP and Kaplan NO (Eds). Methods in enzymology (pp. 149–150). Academic Press, New York.

Bolin, D.W., King, R.P., & Klosterman, E.W. (1952). A simplified method for the determination of chromic oxide (Cr2O3) when used as an index substance. Science, 116(3023), 634–635.

De Silva, S.S., & Anderson, T.A. (1995). Fish Nutrition in Aquaculture (pp.319) Chapman & Hall Aquaculture Series, London.

DeFronzo, R.A., & Goodman, A.M. (1995). Efficacy of metformin in patients with noninsulin-dependent diabetes mellitus, The Multicenter Metformin Study Group. The New England Journal of Medicine, 333, 541–549.

Drew, B.S., Dixon, A.F., & Dixon, J.B. (2007). Obesity management: update on orlistat. Vascular Health and Risk Management, 3(6), 817–821.

Dufrane, D., Van Steenberghe, M., Guiot, Y., Goebbels, R.M., Saliez, A., & Gianello, P. (2006). Streptozotocin-induced diabetes in large animals (pigs/primates): role of GLUT2 transporter and beta-cell plasticity. Journal of Transplantation, 81, 36–45.

Dutta, S., & Ray, S. (2020). Comparative assessment of total phenolic content and in vitro antioxidant activities of bark and leaf methanolic extracts of Manilkara hexandra (Roxb.) Dubard. Journal of King Saud University-Science, 32, 643–647.

Fiske, C.H., & Subbarow, Y. (1925). The colorimetric determination of phosphorus. Journal of Biological Chemistry, 66, 375–400.

Freeland, R.A., & Harper, A.L. (1959). The study of metabolic pathway by means of adaptation. Journal of Biological Chemistry,234, 1350–1354.

Gaur, V., & Mathur, A. (2019). Determination of LC50 of phenolic compounds (phenol & m-cresol) for a fish, labeorohita. International Journal of Medical Laboratory Research, 4(1), 55–63.

Gleeson, M., Connaughton, V., & Arneson, L. (2007). Induction of hyperglycaemia in zebrafish (Danio rerio) leads to morphological changes in the retina. Acta Diabetologica, 44, 157–163.

Hertz, Y., Madar, Z., Hepper, B., & Gertler, A. (1989). Glucose metabolism in the common carp (Cyprinus carpio, L.) the effects of cobalt and chromium. Aquaculture, 76, 255–267.

Hori, T.S.F., Avilez, I.M., Inoue, L.K., & Moraes, G. (2006). Metabolical changes induced by chronic phenol exposure in matrinxãBryconcephalus (Teleostei: Characidae) juveniles. Comparative Biochemistry and Physiology Part C: Toxicology & Pharmacology, 143(1), 67–72.

Kornberg, A., & Horecker, B.L. (1955). Glucose-6-phosphate dehydrogenase. In: Colowick SP and Kaplan NO (Eds) Methods in enzymology (pp. 323–333). Academic Press, New York.

Landin, K., Tengborn, L., & Smith, U. (1991). Treating insulin resistance in hypertension with metformin reduces both blood pressure and metabolic risk factors. Journal of International Medicine, 229, 181–187.

Liao, Y., Takashima, S., Zhao, H., Asano, Y., Shintani, Y., Minamino, T., Kim, J., Fujita, M., Hori, M., & Kitakaze, M. (2006). Control of plasma glucose with alpha-glucosidase inhibitor attenuates oxidative stress and slows the progression of heart failure in mice. Cardiovasc Research, 70, 107–116.

Lowry, O.H., Rosebrough, N.J., Farr, A.L., & Randall, R.J. (1951). Measurement of protein with the Folin phenol reagent. Journal of Biological Chemistry,193, 265–275.

Maitra, S., & Ray, A.K. (2003). Inhibition of digestive enzyme in rohu, Labeo rohita (Hamilton), fingerlings by tannin: an in vitro study. Aquaculture research, 34, 93–95.

Malik, S.K., Choudhary, R., Kumar, S., Dhariwal, O.P., & Deswal, R.P.S. (2012). Socio-economic and horticultural potential of KhirniManilkara hexandra (Roxb.) Dubard: a promising underutilized fruit species of India. Genetic Resources and Crop Evolution, 59, 1255–1265.

Mark, R. O., Elena, D., & Andrew, P.H. (2000). Evidence that metformin exerts its anti-diabetic effects through inhibition of complex 1 of the mitochondrial respiratory chain.Biochemical. Journal, 348, 607–614.

Misra, G., & Mitra, R. (1968). Mimusops hexandra-Iii. Constituents of root, leaves and mesocarp. Phytochemistry, 7, 2173–2176.

Mondal, S., & Ghosh, K. (2018). Utilization of fermented Pistia leaves in the diet of Rohu, Labeo rohita (Hamilton): Effects on growth, digestibility and whole body composition.Waste and Biomass Valorization,10, 3331–3342.

Mondal, S., & Ghosh, K. (2010). Inhibitory effect of Pistia tannin on digestive enzymes of Indian major carps: an in vitro study. Fish Physiology and Biochemistry, 36, 1171–1180.

Monfared, A. L., & Salati, A. P. (2012). Histomorphometric and biochemical studies on the liver of rainbow trout (Oncorhynchus mykiss) after exposure to sublethal concentrations of phenol. Toxicology and Industrial Health, 29(9), 856–861.

Nimbekar, T.P., Katolkar, P.P., & Patil, A.T. (2012). Effects of Manilkara hexandra on blood glucose levels of normal and alloxan induced diabetic rats. Research Journal of Pharmacy and Technology, 5(3), 367–368.

Ogden, C.L., Carroll, M.D., Curtin, L.R., McDowell, M.A., Tabak, C.J., & Flegal, K.M. (2006). Prevalence of overweight and obesity in the United States, 1999-2004. JAMA,295(13), 1549–1555.

Okokon, J.E., Bassey, A. L., & Obot, J. (2006). Antidiabetic activity of ethanolic leaf extract of Croton zambesicus Muell (thunder plant) in alloxan diabetic rats, The African Journal of Traditional, Complementary and Alternative Medicines, 3(2), 21–26.

Olsen, A.S., Sarras, M.P., & Intine, R.V. (2010). Limb regeneration is impaired in an adult zebrafish model of diabetes mellitus. Wound Repair and Regeneration,18(5), 532–542.

Önal, S., Timur, S., Okutucu, B., & Zihnioğlu, F. (2005). Inhibition of α□glucosidase by aqueous extracts of some potent antidiabetic medicinal herbs. Preparative Biochemistry and Biotechnology, 35, 29–36.

Parekh, J. S., & Chanda, A. (2010). Antibacterial activity of aqueous and alcoholic extracts of Indian medicinal plants against some Staphylococcus Species. Turkish Journal of Biology, 32, 63–71.

Patel, E.D., & Patel, N. J. (2015). Antidiabetic activity of leaves of Manilkara hexandra: role of carbohydrate metabolising α-amylase enzyme. Advance research in Pharmaceuticals and Biologicals, 5(2), 863–867.

Patel, M., & Mishra, S. (2011). Hypoglycemic activity of alkaloidal fraction of Tinospora cordifolia. Phytomedicine,18(12), 1045–1052.

Raghavan, B., & Krishna Kumari, S. (2006). Effect of Terminalia arjuna stem bark on antioxidant status in liver and kidney failure of alloxan diabetes rats. Indian Journal of Physiology and Pharmacology, 50(2), 133–142.

Risbud, M.V., & Bhonde, R.R. (2002). Models of pancreatic regeneration in diabetes. Diabetes Research and Clinical Practice,58, 155–165.

Robinson, A. C., Burke, J., Robinson, S., Johnston, D. G., & Elkeles, R.S. (1998). The effects of metformin on glycemic control and serum lipids in insulin-treated NIDDM patients with suboptimal metabolic control. Diabetes Care, 21, 701–705.

Seyedan, A., Alshawsh, M.A., Alshagga M.A., Koosha, S., & Mohamed, Z. (2015). Medicinal plants and their inhibitory activities against pancreatic lipase: A Review. Evidence Based Complementary and Alternative Medicine, 2015, 1–13.

Srinivasan, K., Viswanad, B., Asrat, L., Kaul, C.L., & Ramarao, P. (2005). Combination of high-fat diet-fed and low-dose streptozotocin-treated rat: a model for type 2 diabetes and pharmacological screening. Natural Product Pharmacolog,52, 313–320.

Surwit, R.S., Kuhn, C.M., Cochrane, C., McCubbin, J.A., & Feinglos M.N. (1988). Diet-induced type II diabetes in C57BL/6J mice. Diabetes. 37, 1163–1167.

Tranulis, M.A., Dregni, O., Christophersen, B., Krogdahl, A., & Borrebaek, B. (1996). A glucokinase-like enzyme in the liver of atlantic salmon (Salmo salar). Comparative Biochemistry and Physiology. Part B: Biochemistry & molecular biology, 114, 35–39.

Van Weel, C. (2005). α-glucosidase inhibitors for patients with type 2 diabetes results from a cochrane systematic review and meta-analysis. Diabetes Care, 28, 154–163.

Walter, H.E. (1984). Proteinases: methods with hemoglobin, casein and azocoll as substrates. In: Bergmeyer, H.U. (Eds), Methods of Enzymatic Analysis (pp. 270–277). Verlag Chemie, Weinheim.

Warrier, P.K. (1993). Indian medicinal plants a compendium of 500 species (Vol.5). Orient Blackswan.

Westergaard, N., Brand C. L., & Lewinsky, R. H. (1999). Peroxy vanadium compounds inhibit glucose-6-phosphatase activity and glucagon-stimulated hepatic glucose output in the rat in vivo. Archives of Biochemistry and Biophysics, 366, 55–60.

Xia, X., Yan, J., Shen, Y., Tang, K., Yin, J., Zhang, Y., Yang, D., Liang, H., Ye, J., & Weng, J., (2011). Berberine Improves glucose metabolism in diabetic rats by inhibition of hepatic gluconeogenesis. Plos one, 6(2), 1–10.

Zang, L., Shimada, Y., & Nishimura, N. (2017). Development of a novel zebrafish model for Type 2 diabetes mellitus. Scientific Reports,7, 1461, 1-11.

